# Morphological colour adaptation during development: Involvement of Growth Hormone Receptor 1

**DOI:** 10.1101/2020.06.01.128538

**Authors:** Tomás Horacio Delgadin, Diana Carolina Castañeda-Cortés, Clara Sacks, Andrés Breccia, Juan Ignacio Fernandino, Paula Gabriela Vissio

## Abstract

Morphological background adaptation is both an endocrine and a nervous response, involving changes in the amount and shape of chromatophores. However, if this adaptation takes place at early developmental stages is largely unknown. Somatolactin (SL) is a pituitary hormone present in fish, which has been associated to skin pigmentation. Moreover, growth hormone receptor type 1 (*ghr1*) has been suggested to be the SL receptor and was associated to background adaptation in adults. In this context, the aim of this work was to evaluate the ontogeny of morphological adaptation to background and the participation of *ghr1* in this process. We found in larval stages of *Cichlasoma dimerus* that the number of head melanophores and ir-SL pituitary cells were increased in individuals reared in black backgrounds compared to fish grown in white tanks. In medaka (*Oryzias latipes)* larval stages a similar response was observed that is altered by a *ghr1* biallelic mutations using CRISPR/*cas9.* Interestingly, melanophore and leucophore numbers are highly associated. Furthermore, we found that somatic growth is reduced in *ghr1* biallelic mutant medaka, establishing the dual function of this growth hormone receptor. Taken together, these results show that morphological background adaptation is present at early stages during development and that is dependent upon *ghr1* unless during this period.

## Introduction

Skin pigmentation of vertebrates is mainly due to the presence of neural crest derived cells termed chromatophores, which contains either light absorbing or light reflecting pigments in specialized intracellular structures called chromatosomes [1]. The pattern of pigmentation, as well as the body animal colour displays during its life is highly variable in vertebrates, constituting a good example of phenotypic plasticity [2]. Moreover, these features can change as a result of developmental constraints at embryo/larvae, larvae/juvenile or juvenile/adult transitions, or as a response to biotic or abiotic environmental cues such as nutrition, UV incidence, surrounding luminosity and social interactions [3]. This process, termed morphological colour change, occurs during long periods of time and implies variations of chromatophore number or density and/or modifications in the chromatophore structure, such as cell size or pigment content [4]. Another related process animals can display is physiological colour change, which occurs in short periods of time as an immediate response to environment changes by aggregating or dispersing chromatophore pigments inside the cell [1]. Both morphological and physiological colour changes are processes driven by endocrine and nervous systems.

Background adaptation, an example of physiological colour change, is widely observed in animals and refers to the organism ability to change its body colour and/or its pattern of pigmentation as a consequence of changes in surrounding luminosity, as for instance dark or bright backgrounds. Interestingly, if background adaptation in adult fishes takes place during long periods of time the physiological colour adaptation can derive into a morphological one [2,3,4]. In this sense, most long term background adaptation studies have analyzed melanophores, a type of chromatophore with a stellate shape containing a black/brown pigment called melanin [1,3,4]. However, it would be interestingly to analyze how chromatophores other than melanophores behaves after long term background adaptation periods, particularly xanthophores, which are light absorbing chromatophore frequently found on fishes, smaller in size than melanophores and contains a yellow/orange pigment [1,5] and leucophores, uncommon light reflective chromatophores characterized by a white appearance. At the same time, the amount of chromatophores in the skin at a given time results from the balance between chromatophore production by differentiation and proliferation of stem cell and removal by apoptosis [4,6]. Finally, as far as we are concern, it is not known if such background adaptation process that is observed in adults could take place during early developmental stages. In this sense, it has been postulated that endocrine and nervous systems are not functional at embryo stages and that the genetic control by which chromatophores are produced in these stages is different than those occurring during morphological colour adaptation in adults [4].

Somatolactin is a fish pituitary hormone, that along with growth hormone (GH) and prolactin (PRL) form a family of pituitary hormones, which are similar in structure, function, and gene organization [7]. Several works suggested a role of this hormone in a number of processes such as reproduction, stress responses, Ca2 + homeostasis, acid-base balance, growth, metabolism, immune responses and skin pigmentation [8, 9, 10, 11, 12, 13, 14, 15, 16, 17,18, 19, 20]. In relation to this, it has been proposed that SL is involved in the generation of chromatophores and regulation of pigment movements in them [18,19,21]. In red drum *Sciaenops ocellatus* and *C. dimerus*, SL was increased in pituitaries of adult fish adapted to dark backgrounds [18,19], and in red drum it was concomitant with an increase in plasma SL protein concentration[19]. Moreover, the mRNA of the putative SL receptor has been localized in epidermis and dermis cells from fish scales and has shown changes associated to SL-likely target cells (melanophores) [20]. What is more, in medaka SL mutant (*ci* strain) defects in chromatophore proliferation and morphogenesis was observed [21], and the transgenic over expression of SL into de *ci* genome rescued the wild phenotype [22]. Nevertheless, the evidence accumulated over the years has not completely unraveled the identity of the SL receptor (SLR) in fish. Early studies carried out in salmonids have concluded that *ghr1* is actually the SLR [23] and *ghr2* the GH receptor [24]. Although studies in other fish species give support for this hypothesis [20,25], recent research carried out in zebrafish has concluded that SL is not a ligand for *ghr1*, moreover, GH is a ligand for both *ghr1* and *ghr2* [26]. Thus, additional work is needed to explore the role of SL and *ghr1* on background adaptation process and somatic growth in fish.

In this study, we provide evidence for the role of SL and *ghr1* in the regulation of background adaptation during the early stage of development in two fish species, *C. dimerus* and medaka *O. latipes*. Additionally, since *ghr1* and *ghr2* are paralogs, we demonstrate that *ghr1*, in addition to presenting a role in background adaptation, also has a role in somatic growth.

## Results

### C. dimerus

#### Background colour effect on melanophore number

First, we examined the onset of background colour adaptation during development as the moment when the fish is first able to sense and respond to different backgrounds. Thus, we started the background colour adaptation experiment right after egg fertilization. From 15 days post hatching (dph), number of melanophores was already higher in larvae raised in black than in white tanks (W: 185±11 vs B: 248±23; p=0.032) and difference was increased at 21 dph (W: 188±11 vs B:264±21; p=0.014) and 30 dph (W:209±8 vs B:327±27; p<0.001; Figure 1b,c,d). No differences were observed at 10 dph (W:192±11 vs B:216±16; p>0.05; Figure 1d). Additionally, standard length was not different between treatments at any time (p>0.05; Figure 1e).

**Figure 1.**
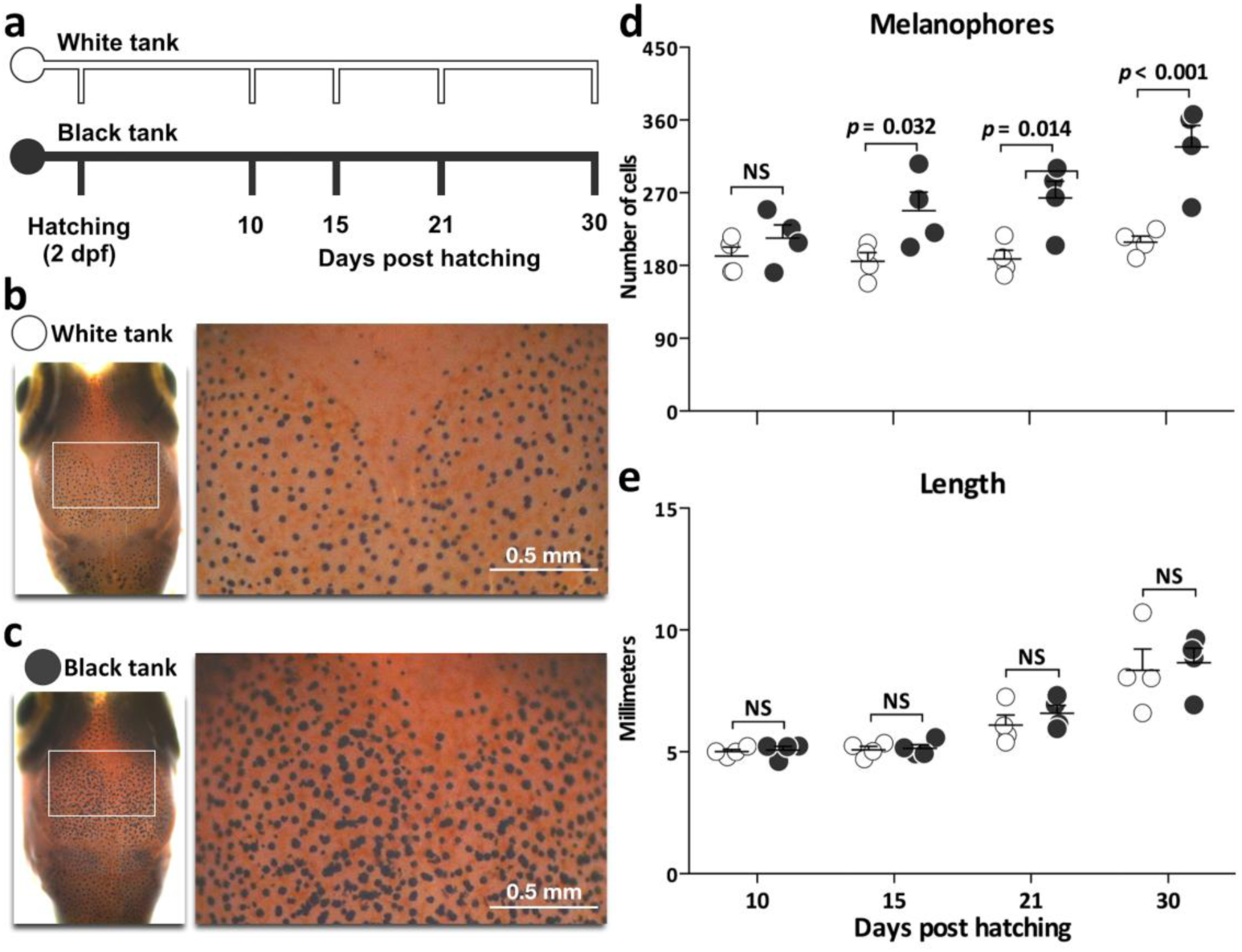
Background adaptation of *C. dimerus*. **a.** Schematic representation of experimental design; embryos exposed since the first day of fertilization (dpf) until 30 dph to white and black tank environments. The collection of samples was at 10, 15, 21 and 30 dph. **b.** White tank larvae at 30 dph. **c.** Black tank larvae at 30 dph. **d.** Number of melanophores in the head of individuals reared in white (white dots) and black (black dots) tanks at different dph. **e.** Standard length of individuals to each treatment at different dph. In **c** and **d**, each point represents the mean value of three larvae.

#### Background colour effect on ir-SL cells

As background colour adaptation was present since 15 dph, we asked if these changes in melanophore number of white and black backgrounds adapted larvae could be related to changes in the pituitary SL cells. The immunohistochemistry assays revealed a higher number of ir-SL cells in black adapted larvae at 30 dph (Figure 2a,b,c; p=0.009). Both cell and nuclear size of ir-SL cells were higher in larvae from black backgrounds (p=0.015 and p=0.004 respectively; Figure 2a,b,d,e).

**Figure 2.**
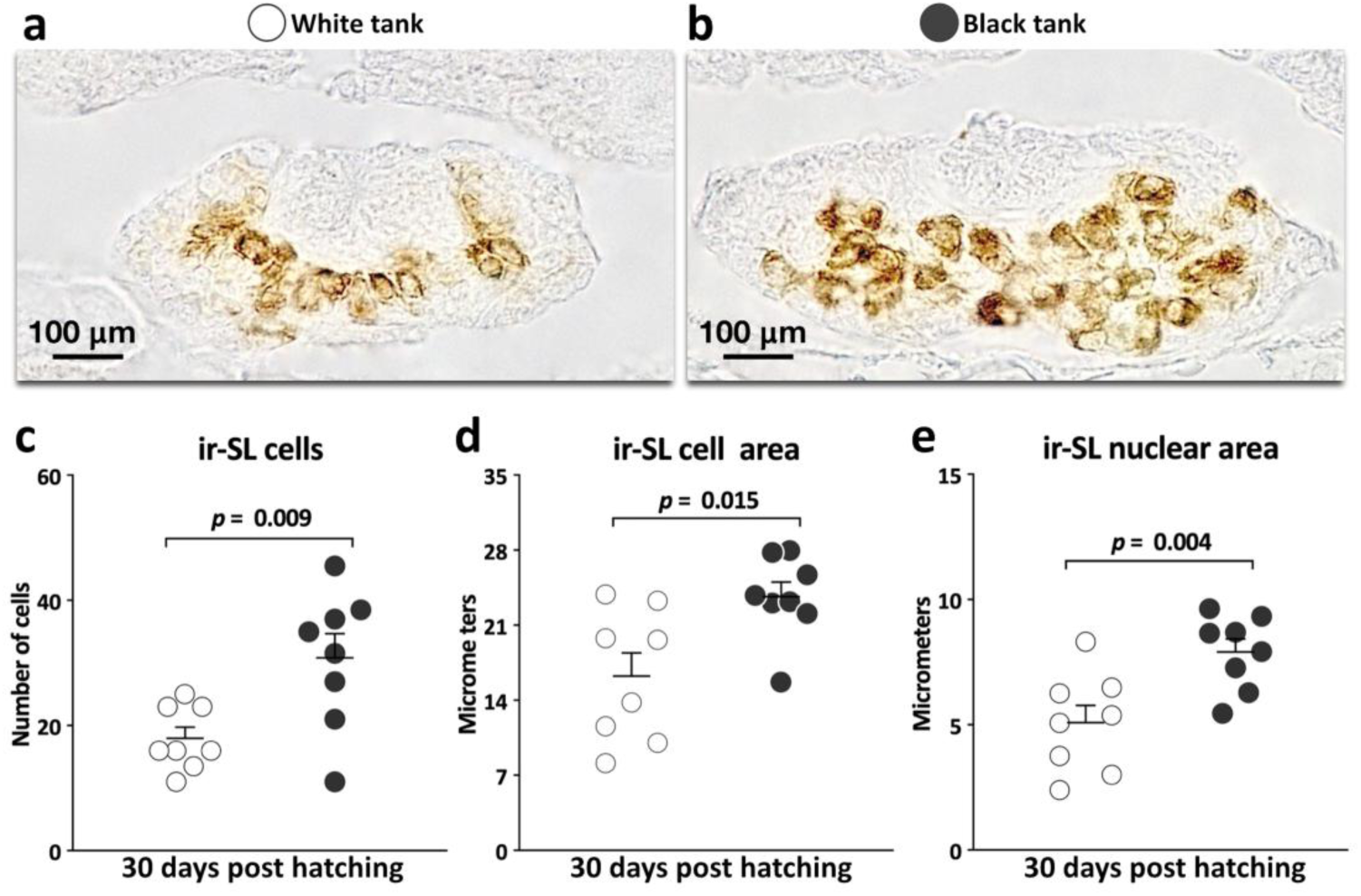
Somatolactin response to background adaptation of *C. dimerus*. Transversal sections of the pituitary gland with ir-Sl cells positive of individuals exposed to white **(a)** and black **(b)** tank environments since the first day of fertilization until 30 days post hatching. Number of ir-Sl cells **(c)**, quantification of ir-Sl cell area **(d)** and ir-Sl nuclear area **(e)** of individuals reared in white tank (white dots) and black tank (black dots) at 30 dph. Sample size by treatment n=8.

### Medaka

#### Background colour effect on chromatophores number

To test if *ghr1*, the SL putative receptor, is involved in colour background adaptation we used medaka fish. Thus, we first had to evaluate if medaka, like *C. dimerus*, is able to adapt to white and black backgrounds from early stages of development (Figure 3a). Medaka larvae raised in black tanks presented a higher number of melanophores than white raised ones at 12 dph (W: 46±3 vs B: 63±4; p=0.016) and 16 dph (W: 38±4 vs B: 96±12; p<0.001) while no differences were observed at 9 dph (Figure 3b,c,d). Standard length was no different between both treatments (p>0.05; Figure 3e).

**Figure 3.**
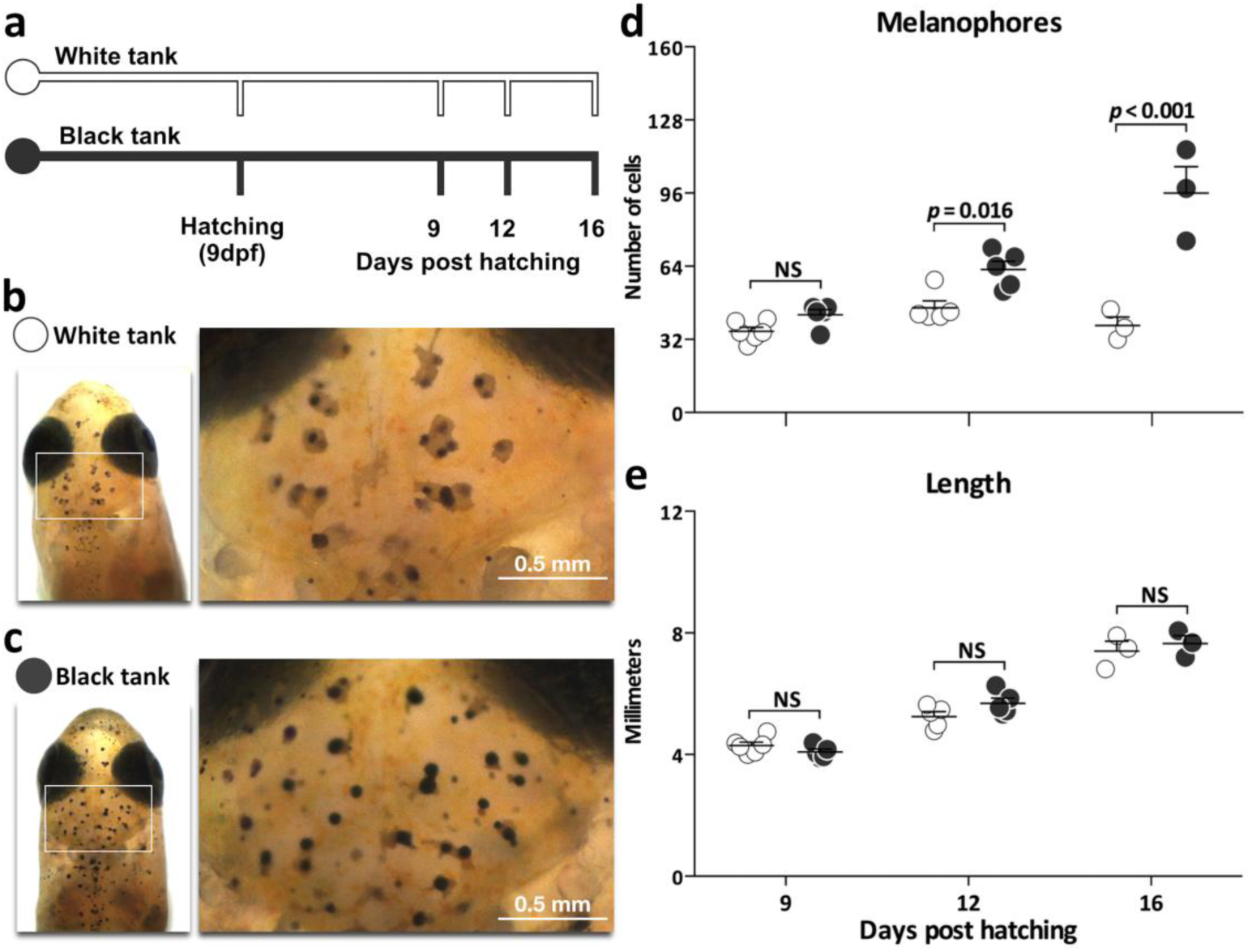
Background adaptation of *O. latipes*. **a.** Schematic representation of experimental design; embryos exposed since the first dpf until 16 dph to white and black tank environments. Samples were collected at 9, 12, and 16 dph. **b.** White tank larvae at 16 dph. **c.** Black tank larvae at 16 dph. **d.** Number of melanophores in the head of individuals from white tank (white dots) and black tank (black dots) at different dph. **e.** Standard length of individuals to each treatment at different dph. Simple size by treatment ranged between n=3 to n=6.

#### Background colour effect on melanophore, leucophore and xanthophore number in ghr1 biallelic mutant larvae

Once we corroborated the background colour adaptation in medaka larvae, our next step was to analyze the participation of *ghr1* in this process generating biallelic mutations of this gene (Figure 4). We obtained F0 individuals with biallelic mutations on *ghr1* (also known as crispants) with a 100% efficiency of injected eggs as determined by heteroduplex mobility assay (HMA; Figure 4b). We confirmed the presence of indels on exon 2 of *ghr1* (Figure 4a) by the presence of multiple bands of heteroduplex (Figure 4b), and confirmed later by sequencing (Figure 4c). Additionally, no indels on potential off-target sites were observed, as can be seen by the presence of a single band in the HMA assay (Figure 4b).

**Figure 4.**
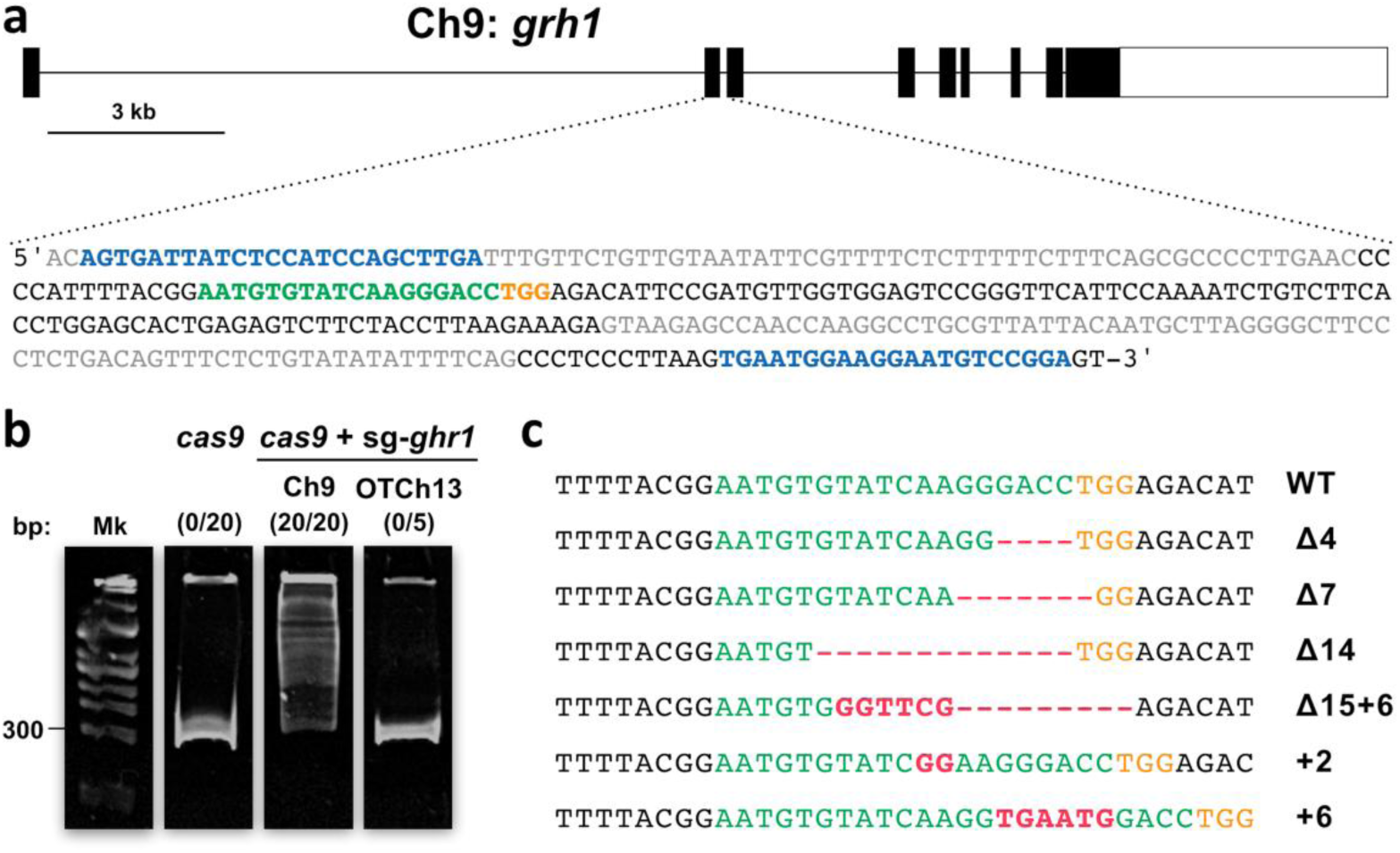
Biallelic mutations of *ghr1* in *O. latipes* using CRISPR/Cas9. **a.** Schematic representation of the genomic structure of grh1 (ENSORLG00000004053). Coding exon regions are shown as solid boxes. The targeting sequence of sgRNA for *ghr1* is indicated by a gray box, adjacent to NGG protospacer adjacent motif (PAM) sequence in black box. **b.** Images of heteroduplex mobility assay (HMA) to analyze the efficiency of the CRISPR/cas9 system and detection of off-target alterations. Representative HMA are shown to embryos microinjected with only cas9 and embryos microinjected with cas9+sg-grh1: amplification to target gene *ghr1* in chromosome 9 (ch9) and potential off-target loci in chromosome 13 (OTCh13) (full-length gels are presented in Supplementary Figure S1). Number of embryos with biallelic mutations/total of eggs injected is shown in parentheses. **c.** Subcloned sequences observed in the cas9 (WT) and cas9+sg-*ghr1* embryos at F1. Red dashes and letters indicate the identified mutations. The size of indels is shown to the right of each mutated sequence.

Since *ghr1* is a paralog of *ghr2*, well demonstrated that it participates in somatic growth, we analyzed firstly the standard length of biallelic *ghr1* mutant larvae raised in white and black backgrounds (Figure 5a). We observed that biallelic *ghr1* mutants showed a decrease in somatic growth compared to wild type larvae, independently of the colour of background (Figure 5b). Additionally, we observed no differences in chromatophore colour between treatments and strains. Visualization with incident light shows black melanophores, orange-creamed leucophores and orange xanthophores, while observation with transmitted light shows same appearance for melanophores and xanthophores but a different colour (brownish) for leucophores (Figure 5c,d,e,f).

**Figure 5.**
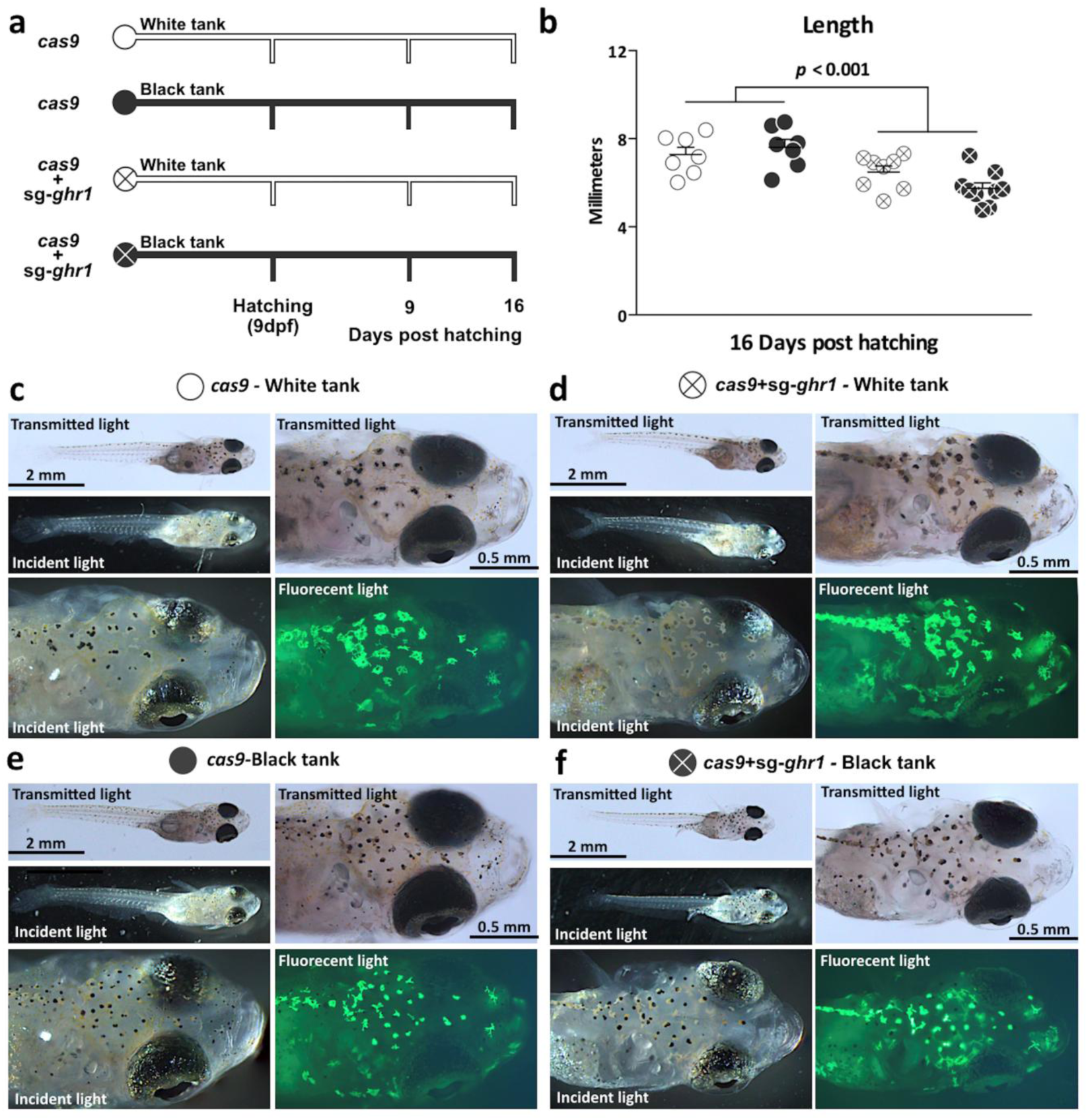
Background adaptation of individuals with biallelic mutations of grh1. **a.** Schematic representation of experimental design; embryos microinjected with cas9 or cas9+sg-grh1 were expose since fertilization until 16 dph to white and black tank environments. Samples were collected at 9, 12, and 16 dph. **b.** Standard length of individuals from each treatment: White tank: cas9 (white filled dots, n=7), cas9+sg-grh1 (crossed out white dots, n=8) and Black tank: cas9 (black filled dots, n=7), cas9+sg-grh1 (crossed out black dots, n=8) at 16 dph. Photomicrographs with different light point exposure and wavelength to observe the types of chromatophores in individuals exposed to white tank: cas9 (white filled dots) (c), cas9+sg-grh1 (crossed out white dots) (d) and Black tank: cas9 (black filled dots) (e), cas9+sg-grh1 (crossed out black dots) (f) at 16 dph.

Then, we submitted biallelic *ghr1* mutated medaka larvae to background colour adaptation experiments to evaluate the involvement of *ghr1* on this process (Figure 5a). No differences were found between mutants and wild type medakas at 9 dph in response to black and white backgrounds (Figure 6a,b,c; p_strainXbackground_>0.05). However, differences were observed at 16 dph, where the interaction effect between strain and background was significant for all chromatophores (p_strainXbackground_ <0.001; Figure 6a,b,c). Specifically, colour background adaptation was observed in control *cas9* medaka (strain injected only with *cas9*), in which black adapted ones presented more melanophores and leucophores than white adapted ones (p<0.001; Figure 6a,b). On the contrary, *ghr1* biallelic mutants failed to adapt to the background as melanophore and leucophore numbers between black and white treatments were not different (p>0.05; Figure 6a,b).

**Figure 6.**
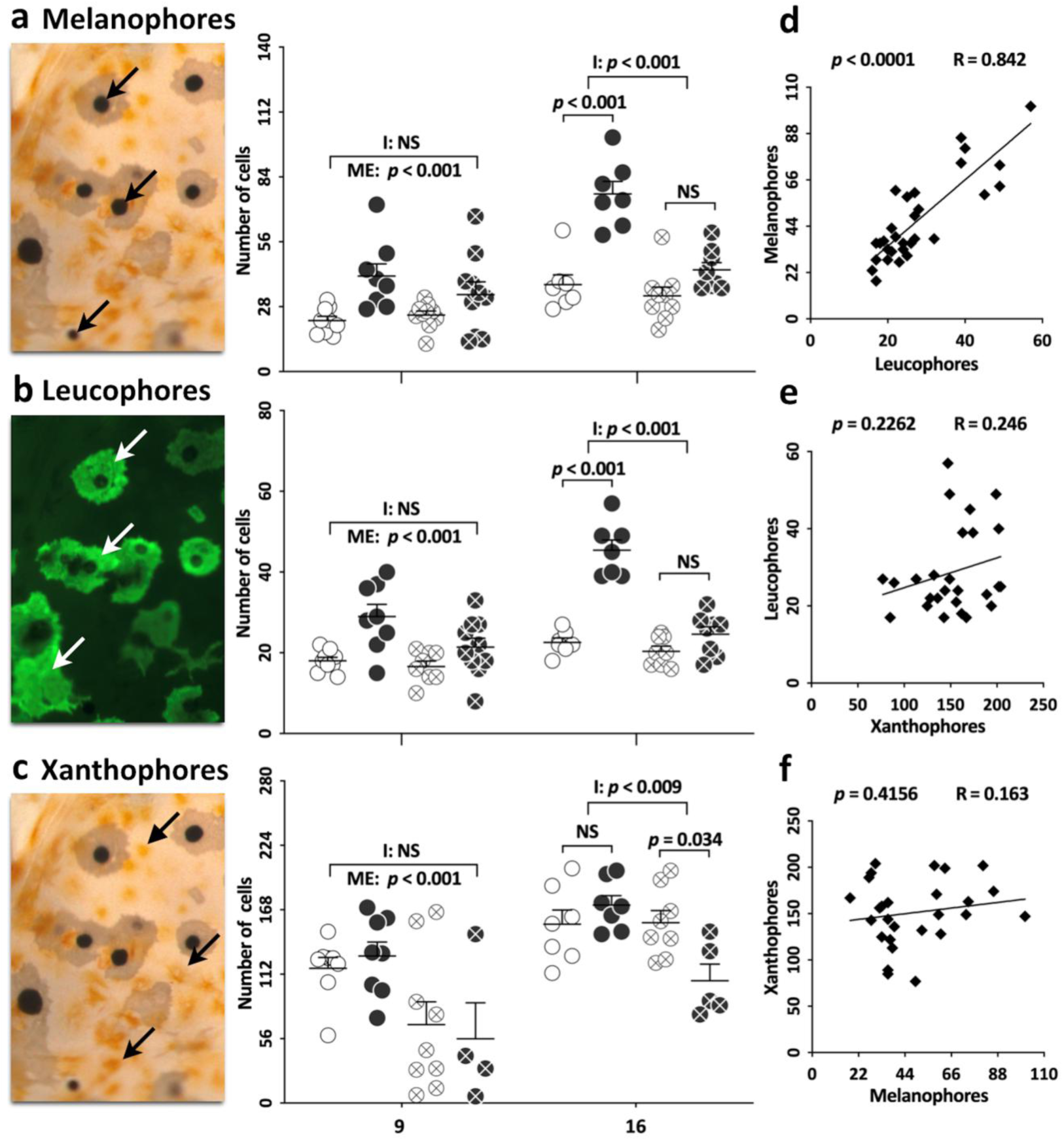
Number of chromatophores in individuals with biallelic mutations under background adaptation. Photomicrographs (left panel, arrows indicate each type of chromatophore) and number of (a) melanophores (n=7-10), (b) leucophores (n=7-10) and (c) xanthophores (n=7-10, except for cas9+sg-grh1 in black tank where n=4-5) in the head of individuals reared in white tank: cas9 (white filled dots), cas9+sg-grh1 (crossed out white dots) and black tank: cas9 (black filled dots), cas9+sg-grh1 (crossed out black dots) at 9 and 16 dph. Pearson’s correlation coefficients between melanophores and leucophores (d), leucophores and xanthophores (e), xanthophores and melanophores (f).

On the other hand, the number of xanthophores were not different between black and white adapted fish at 9 dph (Figure 6c; background effect: p>0.05) for both *ghr1* biallelic mutants and *cas9* injected strains (p_strainXbackground_ >0.05), although *ghr1* mutated medaka presented a lower quantity of xanthophores (strain effect: p<0.001). Afterward, at 16 dph there is a background*line interaction effect (p_strainXbackground_ <0.009) where black adapted medaka in biallelic *ghr1* mutated individuals presented a lower number of xanthophores than white adapted ones (p=0.034).

Additionally, we detected a strong positive correlation between melanophore and leucophore number (r=0.84; p<0.0001; Figure 6d). No association between xanthophores and melanophores or leucophores were found (r=0.16, p=0.4 and r=0.25, p=0.22 respectively). It is worth to mention that 9 dph medaka larvae reared in black backgrounds, unlike those from experiment shown on Figure 3, presented more melanophores and leucophores than white adapted fish (main effect p<0.001; Figure 6a,b) both in biallelic *ghr1* mutated fish and *cas9* injected medaka (interaction effect p>0.05).

Lastly, we analyzed the correlation between total length and the number of different chromatophores. Weak correlations of length with melanophores and leucophores but not with xanthophores were observed (Supplementary Figure S2).

It is worthy to note that the lack of background adaptation in *ghr1* mutated medaka can be arguably nothing but only a side effect of its reduced somatic growth (Figure 5b). To evaluate this hypothesis we inspected in detail data from growth and melanophore number from the experiment in wild type medaka (Figure 3) and compared it to *ghr1* mutated medaka (Figure 6). As stated before, lack of background adaptation is observed at 16dph in mutated *ghr1* medaka. At this time, these fish are 6mm long and showed no statistical differences in the number of leucophores and melanophores number. On the other hand, equivalent wild type sized medaka (5.5 mm long) is observed at 12 dph, where black adapted ones presented more melanophores than white adapted fish (Figure 3). These data show that, once the size effect on melanophore number is discarded, only *ghr1* mutated medaka is unable to adapt to black backgrounds.

#### Background colour effect on ir-SL cells in ghr1 biallelic mutant larvae

We finally asked if lack of background colour adaptation seen at 16 dph in *ghr*1 biallelic mutant medaka was related to SL production. Thus, we carried out immunohistochemistry assays on 16 dph larvae (Figure 7a,b,c,d). We found no effect between *ghr1* mutants and *cas9* injected medaka strains in the number of ir-SL cells, their nuclear and cell size (p_strainXbackground_ >0.05; Figure 5e,f,g). However, as previously observed in *C. dimerus*, black adapted medaka from both lines presented more SL-ir cells (p=0.008) with a larger cell and nuclear size (p<0.001) than white adapted ones (Figure 7e,f,g).

**Figure 7.**
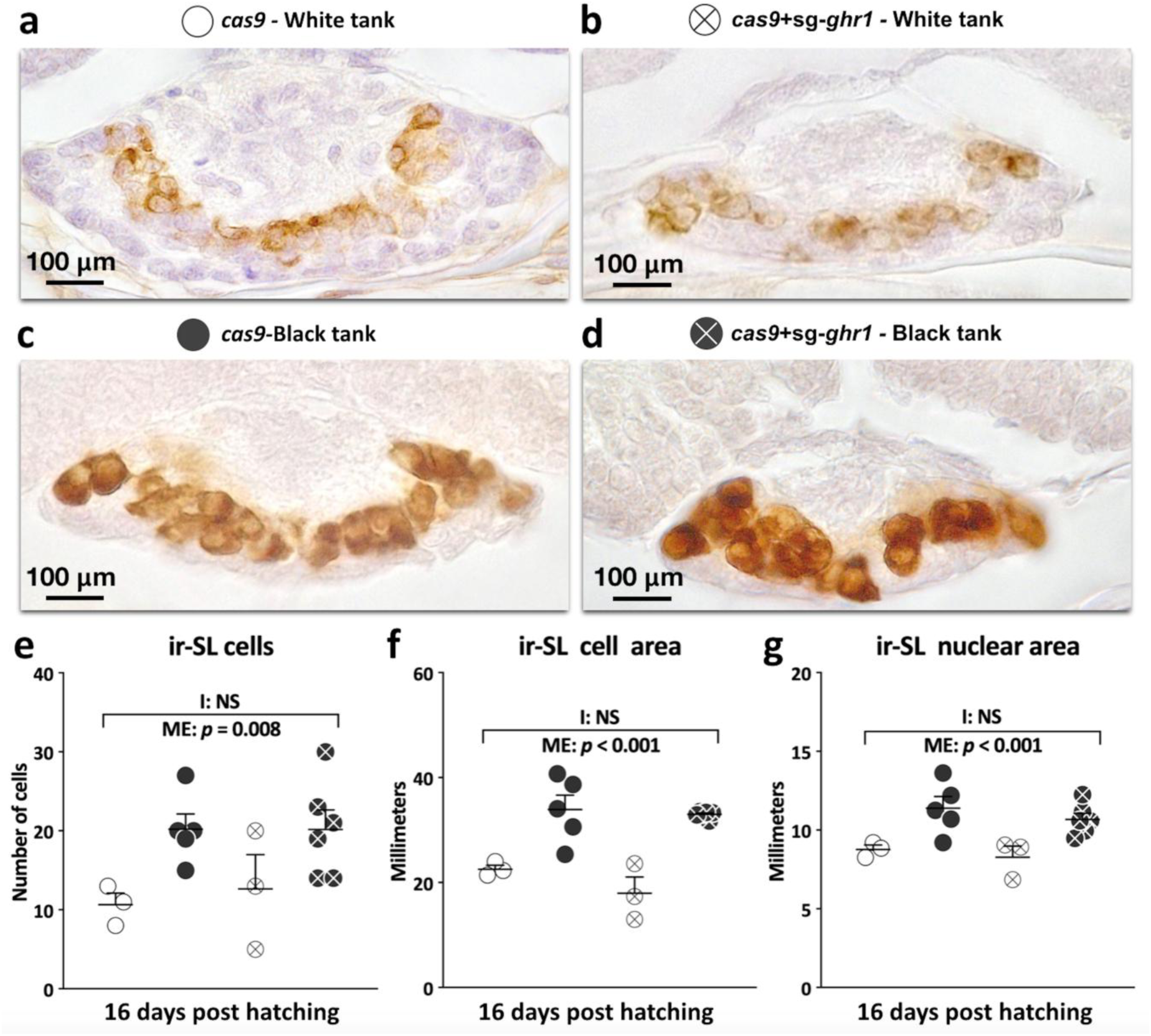
Somatolactin response to background adaptation of individuals with biallelic mutations of *grh1*. Transversal sections of the pituitary gland with ir-Sl cells of individuals exposed to white tank: *cas9* (white filled dots, n=3) **(a)**, *cas9*+*sg-grh1* (crossed out white dots, n=5) **(b)** and Black tank: *cas9* (black filled dots, n=3)**(c)**, *cas9*+*sg-grh1* (crossed out black dots, n=6) **(d)** at 16 dph. Number of ir-Sl cells **(e)**, ir-Sl cell area **(f)** and ir-Sl nuclear area **(g)** of individuals from each treatment at 16 dph.

#### Background colour effect on ir-GH cells and GH receptors in ghr1 biallelic mutant larvae

Given the high structural similarity and phylogenetic relationship between SL and GH, we next evaluated the number of ir-GH cells in the pituitary gland of 16dph medaka (Figure 8c,d,e,f). No differences were observed between the amount of ir-GH cells (Figure 8g), ir-GH cell area (Figure 8h) and ir-GH nuclear area (Figure 8i) in fish reared in black and white and between fish strains.

**Figure 8.**
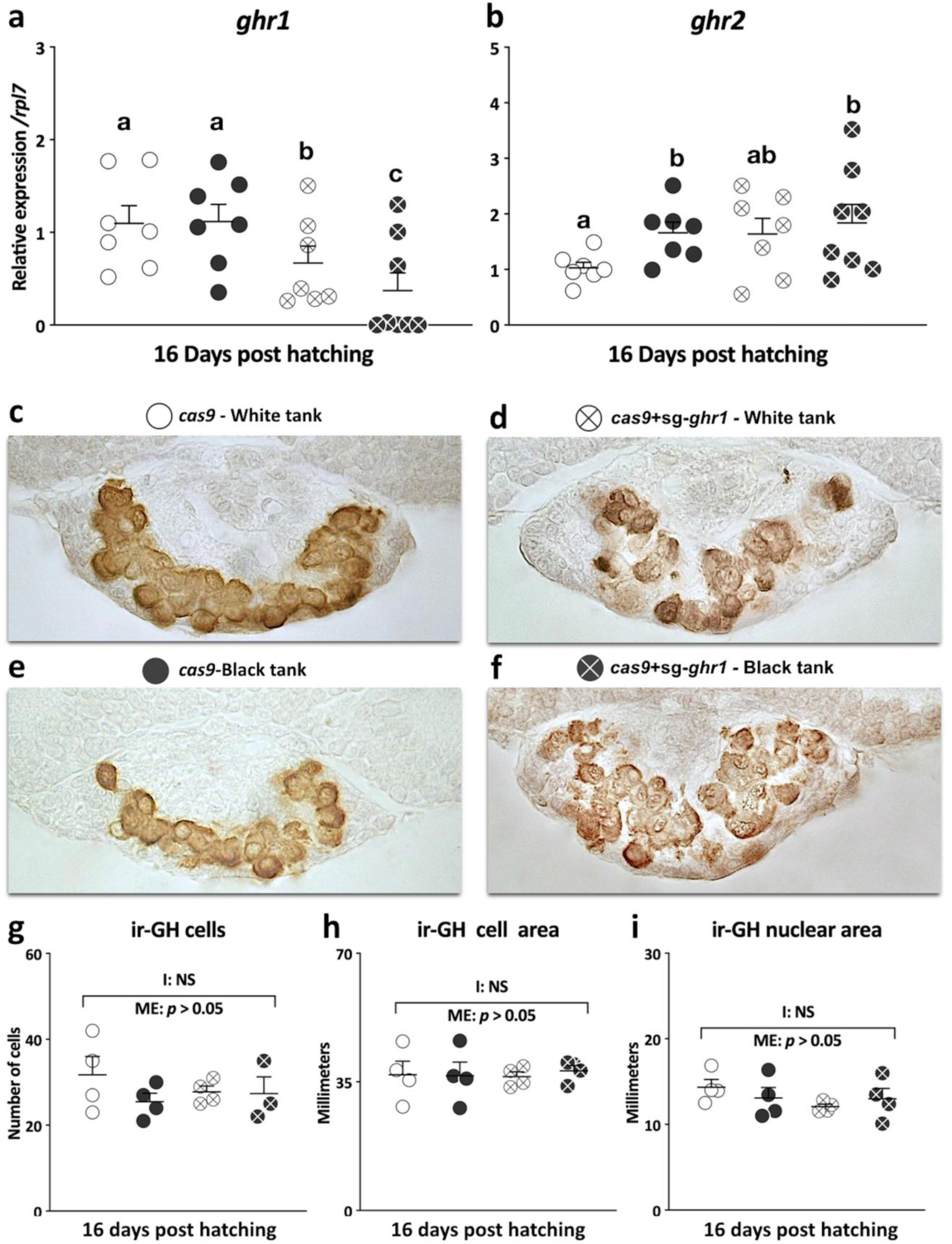
Relative expression of growth hormone receptors and growth hormone response to background adaptation of individuals with biallelic mutations on grh1. Transcript abundance profiles of *ghr1* (a) and *ghr2* (b) from individuals reared in white tank: cas9 (white filled dots), cas9+sg-grh1 (crossed out white dots) and Black tank: cas9 (black filled dots), cas9+sg-grh1 (crossed out black dots) at 16 days post hatching (dph). Gene expression levels are expressed relative to the white tank cas9 (white filled dots) group and normalized to rpl7. Different letters indicate statistically significant differences between treatment. Transversal sections of the pituitary gland with ir-GH cells from individuals exposed to white tank: cas9 (white filled dots) (c), cas9+sg-grh1 (crossed out white dots) (d) and Black tank: cas9 (black filled dots) (e), cas9+sg-grh1 (crossed out black dots) (f) at 16 dph. Number of ir-GH cells (g), ir-GH cell area (h) and ir-GH nuclear area (i) in individuals from each treatment at 16 dph.

Additionally, *ghr1* mRNA levels were lower in *ghr1* mutant medaka (p=0.003), with no differences between individuals from black and white treatments in *cas9* strain (Figure 8a). On the other hand, *ghr2* mRNA expression was higher in black adapted fish for cas9 strain (p=0.005), while this difference was not detected in *ghr2* mutated fish (p=0.326; Figure 8b).

## Discussion

Background adaptation has been widely studied in fish as well as in many other vertebrate species [2,3,4,5]. Nevertheless, most studies were focused on physiological other than morphological changes, that is to say, changes that affect pigment aggregation or dispersion that occurs over short periods of time but not variations in the number of chromatophores on the skin, a process that involves both cell differentiation and proliferation and is referred to as morphological adaptation [4,6]. Although several endocrine and nervous mechanisms have been proposed to regulate morphological adaptation in adult fish, including a central role of SL [21], the onset of such a process during development had remained unexplored until now. Here, we present clear evidence on the occurrence of morphological background adaptation at early developmental stages in two fish species concomitant with variations in SL. Moreover, as far as we are concerned, this is the first work that analyzes the mutation of *ghr1*, a SL putative receptor, demonstrating that morphological background adaptation in the first stages of development is dependent upon this GH receptor. Besides, evidences for a crosstalk between GH and SL to *ghr1 in vivo* are presented.

We first aimed to determine when the fish is able to adapt to the background. Results from *C. dimerus* indicate that the mechanisms by which adaptation takes place must be present early in larvae stages. By this time, 15 dph, the hypothalamic neuropeptide melanin-concentrating hormone (MCH), the *pars* intermedia pituitary hormones melanocyte-stimulating hormone (MSH) and SL are already expressed in *C. dimerus* [27,28], hormones that have been demonstrated to regulate both morphological and physiological changes in adult fish [2,4,18,20,29]. This time co-occurrence between background adaptation and the presence of these hormones shows that skin pigmentation regulating mechanisms are active since early stage of development. A previous study carried out in rainbow trout have shown that physiological background adaptation is present at early stages in ontogeny and that is dependent upon sympathetic innervations, but not dependent on MCH or MSH, both factors present before adaptation took place [30]. This conclusion was based on the fact that physiological background adaptation occurred at 10 dph, but both MCH and MSH expression was not altered by black or white background until 28dph. Unfortunately, the authors did not analyze if morphological background adaptation occurred after 28dph. Although we did not evaluate MCH and MSH in this work, we detected an increment on SL producing cells along with a higher nuclear and cell size in individuals reared in black compared to white backgrounds. Even though we were not able to measure SL levels in plasma, these facts are indicative of an increment of SL in the organism. Moreover, the amount of GH producing cells, as well as its cell and nuclear area were not affected by background or medaka strain, which suggest that GH, unlike SL, is not involved in morphological background adaptation. Together, the increment in melanophores at 15dph on black adapted fish agrees with the hypothesis that SL is involved in background colour adaptation as occurs in adult *C. dimerus* [18, 20].

The function of *ghr1* is not completely understood in fish. As stated above, it was first described as a SL receptor in salmonids, but later *in vitro* studies in other fish species demonstrated that *ghr1* respond to GH, but not to SL [26]. These contradictory results can be attributable to species specific differences owing to divergence and subfunctionalization. Thus, we hypothesized that if *ghr1* is a SL receptor, as proposed for salmonids [23], medaka and other fish species [20,25,31] and considering that *ci* medaka, which lacks of a functional α-SL, presents alterations in chromatophore proliferation and morphogenesis, a *ghr1* mutation in fish should result in affected background adaptation and/or chromatophore production. Therefore, we generated CRISPR/Cas9 mutant for *ghr1* in medaka, where unlike *C. dimerus*, this technique is available. Our results showed that wild type medaka is able to adapt to black backgrounds at 16dph, which indicates that the processes that allow for background adaptation is present before 16dph and after 9dph. Interestingly, morphological adaptation to black background is severely diminished for melanophores and completely abolished for leucophores in *ghr1* deficient medaka when compared to control/*cas9* ones at 16dph. Thus, the lack of morphological adaptation in *ghr1* deficient medaka is in agreement with the hypothesis by which *ghr1* is the SL receptor. This fact does not exclude the possibility that *ghr2*, the *ghr1* paralog, can mediate other biological function of SL. Indeed, we did not observe an effect of *ghr1* mutations on SL producing cells, which may suggest that SL is binding to other receptor as well or an absence of negative feedback loop to this endocrine pathway. In relation to that, we observed that *ghr2* but not *ghr1* is upregulated in black adapted medaka for the control cas9 strain, but it disappears in *ghr1* biallelic mutants. Nevertheless, the no effect of background in *ghr1* expression on cas9 medaka larvae may be due to technical issues, as in this work we started from RNA extracted from whole head, whereas those studies where *ghr1* was upregulated were conducted in isolated scales from adult [20]. Interestingly, *ghr1* mRNA expression is decreased in *ghr1* mutant medaka.

To our knowledge, this is the first study to report the functional role of *ghr1* on background adaptation. However, the role of SL on body pigmentation has been studied on *ci* medaka, a mutant strain which lacks of a functional SL. These *ci* fish showed phenotypic alterations related to proliferation and differentiation of chromatophores [21], particularly they present a grey body colour caused by an increase in the number and size of white leucophores and a decrease in the number of orange xanthophores, with no differences in melanophore number. If *ghr1* was the SL receptor, we would expect that the *ci* and *ghr1* mutant medaka phenotypes were similar. In fact, this is the case except for leucophores. Our results show that *ghr1* mutant medaka contain less xanthophores than control individuals, no variation in melanophores and unlike *ci* medaka, no variation on leucophores number. Interestingly, colour of leucophores on *ghr1* mutants is orange-creamed, as occurs in control individuals, while in *ci* medaka they are white. If *ghr1* was the SL receptor, a possible explanation for this discrepancy could be attributable to the methodology of mutant strain generation. The genetic background of *ci* medaka line may be very different from the strain used in these studies, where *ghr1* mutant medaka was designed specifically for the *ghr1* gene.

On the other hand, as *ghr1* is a GH receptor paralog, it is expected that this receptor maintains functions related to GH function such as growth. Indeed, we showed that somatic growth is reduced in *ghr1* mutated medaka. This result along with the lack of morphological background adaptation in this *ghr1* medaka strain strongly suggests a crosstalk between GH and SL to *ghr1 in vivo*. This crosstalk is in agreement with studies carried out in *ci* medaka that overexpresses GH ectopically, where this fish showed a higher body size, as expected, and surprisingly a higher number of melanophore in the skin than control fish [32], although in a lesser extent than *ci* medaka that overexpresses SL ectopically [22].

Finally, it should be mentioned that although *ghr1* regulates morphological background adaptation it does not affect normal melanophore and leucophore production as larvae present a normal colour appearance with no evident difference in the amount of these chromatophore between *ghr1* mutated and cas9 medaka. Thus, how is it possible that *ghr1* is able to regulate morphological adaptation without affecting the production of chormatophores? A potential explanation would be that *ghr1* is involved in chromatophore survival. A number of studies on mammalian models have demonstrated survival and protective effects of GH in immune cells through *ghr* [33], and as *ghr1* is a GH receptor paralog, this function may still remain for *ghr1*. As stated before, the number of chromatophores in the skin is a result of production by proliferation/differentiation and elimination by apoptosis [4,6]. In this context, high levels of SL induced by long term adaptation to black backgrounds would activate *ghr1* and stimulate the chromatophore survival, but when *ghr1* is mutated, this protective effect would be lost and as a consequence the number of chromatophores would not increase: all newly produced chromatophores would be eventually eliminated by apoptosis. Our results showed that black adapted fish present more SL producing cells with higher cell and nuclear size, which indirectly suggest higher levels of SL and gives support to the survival hypothesis. However, a better method of SL quantification should be used to this purpose.

Both *ghr1* mutated medaka and *ci* medaka lack of morphological adaptation for leucophores [21]. In this species it was reported that adaptation to white backgrounds conducts to a higher number of leucophores in the trunk of the body [6], but in *ci* medaka this increment does not take place [21]. In our experiments the number of leucophores increased in wild type black adapted medaka, but this increment was not observed in *ghr1* mutated medaka. Besides the opposite response of leucophore to background adaptation, it is important to note that both *ci* and *ghr1* mutated medaka showed no morphological background adaptation, which suggest that both SL and *ghr1* play a role in leucophore number regulation in long term background adaptation. The opposite response in the number of leucophores in wild type to long term background adaptation could be explained by the region where they were quantified. The experiments carried out by Sugimoto *et al*[6] and those of *ci* medaka were measured in the trunk of the fish in adult specimens while the results presented in this work were quantified as the total amount of chromatophores at larvae stages on the dorsal side of the head. Additionally, head melanophores and leucophores appeared to be very close in space. In this sense, we observed a strong positive correlation in the number of head melanophores and leucophores. This feature along with a similar response to long term background adaptation suggests a common regulatory mechanism dependent upon *ghr1* for melanophores and leucophores.

The early onset of background adaptation is clear both in *C. dimerus* and *O. latipes*, as determined by the higher number of melanophores in black adapted fish. However, the medaka strain in which the experiments were conducted was hi-medaka, a strain known to have a variable amount of amelanic melanophores but normal leucophores and xanthophores. This fact may obscure data interpretation derived from melanophores. For this reason, the involvement of *ghr1* in morphological background adaptation is mainly supported by the lack of leucophore background adaptation in *ghr1* mutated medaka. Nevertheless, the number of melanophores is mirrored by the behavior of leucophore numbers with a high statistical correlation, which in turn, despite the possible presence of amelanic melanophores, validates the involvement *of ghr1* in melanophore background adaptation as well.

In summary, in this work we present clear evidences about the role of *ghr1* in the regulation of morphological colour background adaptation early during development.

## Materials and methods

### Animals

#### Background adaptation: C. dimerus

Eggs from four independent *C. dimerus* spawning were divided each one into two groups and randomly placed either into 2L white or black tanks right after egg fertilization (Figure 1a). At day 10, 15, 21 and 30 post hatching three larvae from each background were withdrawn and placed on a 20µg/ml dopamine solution to induce melanophore aggregation, placed at 0°C to reduce movements an observed under optical magnifier. All melanophores from dorsal region of the head were manually counted from each larva. Total length was measured at each time point using ImageJ^®^ software after appropriate calibration. Additionally, at least two larvae from each background and spawning were placed at 0°C and then fixed in Bouin’s solution for further histological analysis. All experiments comply with the approval of Comisión Institucional para el Cuidado y Uso de Animales de Laboratorio, Facultad de Ciencias Exactas y Naturales, Buenos Aires, Argentina (protocol number 95).

#### Background adaptation: medaka O. latipes

Medaka eggs were taken from 2 to 3 spawning of different male and female pairs for each time point and randomly assigned to black or white backgrounds, reared in Petri dish until hatching and then transferred to tanks of the same background till sampling at 9, 12 and 16 dph (Figure 3a). To measure dorsal head melanophore number and total length the method previously described for *C. dimerus* was employed. The strain hi-medaka (ID: MT835) was supplied from the National BioResource Project (NBRP) Medaka (www.shigen.nig.ac.jp/medaka/). All fish were maintained and fed following standard protocols to medaka [34]. Fish were handled in accordance with the Universities Federation for Animal Welfare Handbook on the Care and Management of Laboratory Animals (www.ufaw.org.uk) and internal institutional regulations.

### SL immunohistochemistry assays

Larvae from *C. dimerus* and medaka sampled at 16 dph and 30 dph were fixed in Bouin’s solution for 12hs in darkness at 4°C, dehydrated in an increasing alcohol/water graded solution, submerged in xylene for 5min and then embedded in Paraplast^®^ (Fisherbrand, Fisher, WA, USA). 7µm transversal sections at pituitary level were xylene treated for 35min twice and then rehydrated in a graded alcohol series until phosphate-buffered saline solution (PBS, pH=7.4). All sections were incubated for 5 min in 0.3% (v/v) H_2_O_2_ to block endogenous peroxidase. After three washes for 5min in PBS, slices were blocked with 5% nonfat dry milk PBS for one hour and then incubated overnight at 4°C with rabbit anti-*Sparus aurata* SL antiserum (1:1500). Afterward, slices were washed three times in PBS for 5min and incubated with biotinylated anti-rabbit IgG (1:500 in PBS, Sigma-Aldrich) at room temperature (RT) for 1 h. Following three PBS washes, the sections were incubated with IgG peroxidase-conjugated streptavidin (dilution 1 : 500 in PBS; Invitrogen) at RT for 1 h. The final step was accomplished by treating sections with 0.3% (w/v) 3,3-diaminobenzidine in Tris buffer (pH 7.6) and 0.02% (v/v) H_2_O_2_. Then, sections were submitted to a slight counterstaining with hematoxylin followed by a rapid dehydration step in a graded alcohol gradient to xylene and mounted in DPX. Slices were observed in a Microphot FX microscope (Nikon, Tokyo, Japan) and photographed if ir-SL cells were present. Antibodies specificities were previously analyzed [27].

### *Biallelic mutations of ghr1 in medaka* using CRISPR/cas9

To analyze the participation of ghr*1* in the background adaptation we performed a biallelic mutation using CRISPR/Cas9 following the protocol described previously [35,36]. Briefly, target sites were designed using the CCTop - CRISPR/Cas9 target online predictor (crispr.cos.uni-heidelberg.de/index.html)[37], which identified sequence 5′ GG-(N18)-NGG3′ in exon 2 of *grh1* (AATGTGTATCAAGGGACC TGG, Figure 4a). Then, to the single guide RNA (sgRNA) synthesis the annealed oligonucleotides was cloning into the sgRNA expression vector pDR274 (Addgene #42250)[38] and synthesized using the MEGAshortscript T7 Transcription Kit (Thermo Fisher Scientific). Additionally, the capped *cas9* RNA was transcribed from pCS2-nCas9n plasmid (Addgene #47929) and synthesized by mMESSAGE mMACHINE SP6 kit (Life Technologies). The synthesized sgRNAs and *cas9* were purified by RNeasy Mini kit purification (QIAGEN). These RNA sequences were diluted to 50 and 200 ng/µL respectively.

Potential off-target sites in the medaka genome for each sgRNAs were searched using the CCTop - CRISPR/Cas9 target online predictor [37]. Only one potential off-target site was identified in the chromosome 13 (ENSORLG00000003362) and analyzed by Heteroduplex mobility assay (HMA) [36] using the primers: FW 5’ AGCTGTGTCAGCCTGTGAAA 3’ and RV 5’ TGAGCGGGGAAAAACATTAC 3’. Electrophoresis performed in 12% acrylamide gel^38^, stained with ethidium bromide for 15 min before examination.

#### Microinjection into medaka embryos and experimental design to background adaptation

Microinjection was performed into fertilized medaka eggs before the first cleavage as described previously[40], using a Nanoject II Auto-Nanoliter Injector (Drummond Scientific) coupled to a stereomicroscope (Olympus). Embryos injected with only *cas9* were used as controls and randomly assigned to black or white backgrounds. For CRISPR/Cas9 system, 4.6 nL of RNA mixture of 25 ng/µl *sgRNA* and 100 ng/µL *cas9* were co-injected into the embryos.

After microinjection, eggs were distributed randomly to black or white backgrounds and sampled at 12 and 16 dph for each treatment. Firstly, for all individuals of each time sampling and treatment the caudal fin was taken for genomic DNA (gDNA) extraction using conventional saline buffer extraction^41^ to corroborate the biallelic mutations generate by CRISPR/Cas9 system using HMA analysis. Conventional PCR analysis was performed with genomic DNA using primers: FW 5’ AGTGATTATCTCCATCCAGCTTGA 3’ and RV 5’ TGAATGGAAGGAATGTCCGGA 3’ (Figure 4a). The embryos that shown formation of heteroduplexes were considered as embryos with biallelic mutations and taked to analysis.

Finally, the screening of indels was performed in F1 fish. Biallelic mutant adult (F0) medaka were mated with wild-type medaka of the Himedaka strain (WT). Genomic DNA was extracted from each F1 embryo for analysis of mutations by HMA, as described previously. Mutant alleles in each embryo were determined by direct sequencing ghr1 gene region.

#### Analysis of biallelic mutation phenotype in to background adaptation

Medaka fish were sampled for chromatophore counting and standard body length measurements as described above. Melanophore and xantophores were visualized on light magnifier while leucophores were detected by fluorescent light excitement. Total chromatophore number was quantified form the dorsal region of the head.

### RNA extraction and quantification by real time PCR (RT-qPCR)

Total RNA was extracted from heads of individual embryos at 16 dph using 350 μl of TRIzol Reagent (Life Technologies), following the manufacturer’s instructions. RNA quantification and purity were determined on NanoDrop 2000 UV-vis spectrophotometer (Thermo Scientific^®^, USA). Samples were DNase I treated (Sigma Aldrich, USA) to avoid genomic DNA contamination starting from 1 µg of total RNA according to manufacturer’s instructions. cDNA synthesis was carried out with M-MLV enzyme (Promega, Madison WI, USA) for 50 min at 37°C followed by 10 min at 70°C using random oligomers (hexamers) as reaction primers in a 10-µl final volume. RT-qPCR reactions were conducted in a 10-µL final volume with 5 µL of 2X FastStar Universal SyBR green Master (Roche, Switzerland), 1.5 µL of forward/reverse primer mix (Table S1; *ghr1*: 250 nM; *ghr2*: 250 nM; rpl7: 100nM), 2.5 µL of cDNA template and 1 µL of water. The amplification protocol consisted of an initial denaturation cycle at 95°C for 10 min, then 40 cycles of denaturation at 95°C for 30 s and annealing/elongation at 60°C for 30 s, followed by a melting curve from 65°C to 95°C to detect possible non specific PCR products. All primers were designed to give an amplicon size between 100 and 130 base pairs. Samples were run in duplicate and no template controls were performed in every run for each primer pair. Raw fluorescence data from qPCR were exported from the Step One Plus Real-Time PCR System (Applied Biosystems) to LinRegPCR software and analyzed to obtain the cycle threshold (CT) and PCR efficiency [42,43]. PCR efficiencies for all primer pair were 90%. The subsequent quantification method was performed by using an efficiency corrected method for relative expression [44] normalizing against rpl7 gene [45].

### Statistical analysis

*For morphological* adaptation experiments in *C. dimerus* both melanophore number and standard body length were analyzed by repeated measured (RM) ANOVAs, while number, cell and nuclear are of ir-SL cells were analyzed by two tailed Student test. Post hoc multiple comparisons by Holm-Sidak test were conducted after RM ANOVAs. Melanophore number and standard length in wild type medaka for morphological adaptation experiment were analyzed by two-way ANOVAs followed by a Tukey’s test for post hoc multiple comparison. Melanophore, leucophore and xanthophore number in morphological background adaption experiments on *ghr1* mutated medaka were analyzed for 16 dph and 21 dph separately. At each time point, a nested ANOVA design was applied considering strains (*ghr1* mutated/*cas9* control), background (black/white) and its interaction as fixed factors, while experiment replication (two replications) as a random factor nested in strain. A Tukey’s post hoc test was applied whenever the interaction effect was statistically significant. ir-SL cell number was analyzed as explained above for *C. dimerus*. Correlation analysis between chromatophores and standard length were conducted by Pearson’s test. Standard length comparison between *ghr1* mutated and *cas9*/control medaka was conducted by a two way ANOVA considering strain and background as factors. Fold change and statistical analysis of RT-qPCR quantifications were performed using FgStatistics interface (http://sites.google.com/site/fgStatistics/), based on the Relative Expression Software Tool (REST) from Pfaffl [44]. All p-values were corrected for multiple comparisons in order to avoid family wise error rate increments. Data are presented as mean ± standard error of the mean (SEM).

## Acknowledgments

We are especially grateful to Dr. Shoji Fukamachi that not only inspired our study through their works but also contributed in this manuscript through fluid mail feedback. We thank to NBRP Medaka (https://shigen.nig.ac.jp/medaka/) for providing hi-medaka (Strain ID: MT835).

## Data availability

The datasets generated during and/or analyzed during the current study are available from the corresponding author on reasonable request.

## Author contributions statement

T.H.D., J.I.F and P.G.V. conceived the experiments. T.H.D. and D.C.C.C. performed most of the experiments. C.S. and A.B. performed immunohistochemistry assays. T.H.D., D.C.C.C., P.G.V. and J.I.F. analyzed the results, wrote and reviewed the manuscript.

## Figures

**Supplementary Figure S1.**
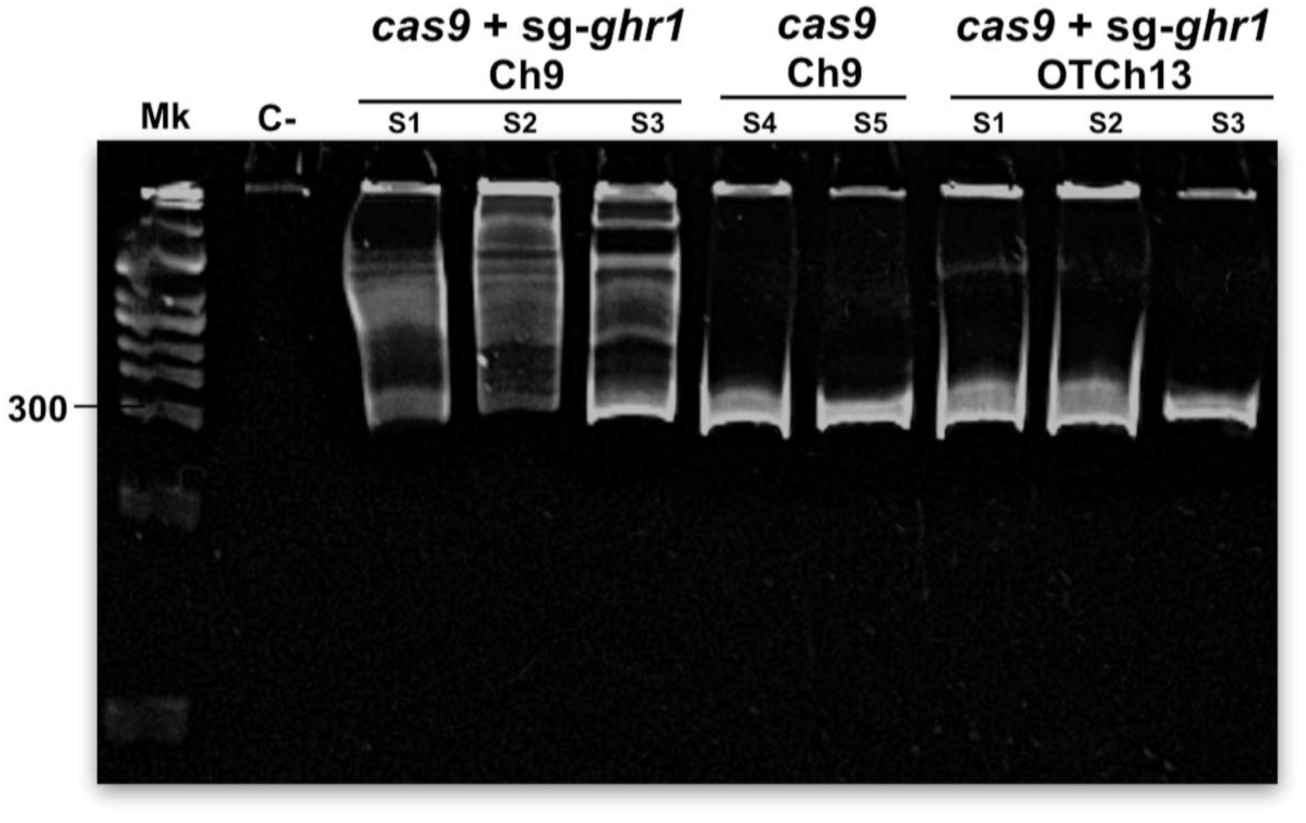
Full-length gel of images of heteroduplex mobility assay (HMA). HMA to analyze to detection of off-target alterations. Representative HMA are show negative control (C-), embryos microinjected with *cas9*+*sg-grh1*: amplification to target gene ghr1 in chromosome 9 (ch9) samples 1,2 and 3 (S1,S2,S3), embryos microinject only with *cas9* samples 4 and 5 (S4,S5) and potential off-target loci in chromosome 13 (OTCh13) in samples 1,2 and 3.

**Supplementary Figure S2.**
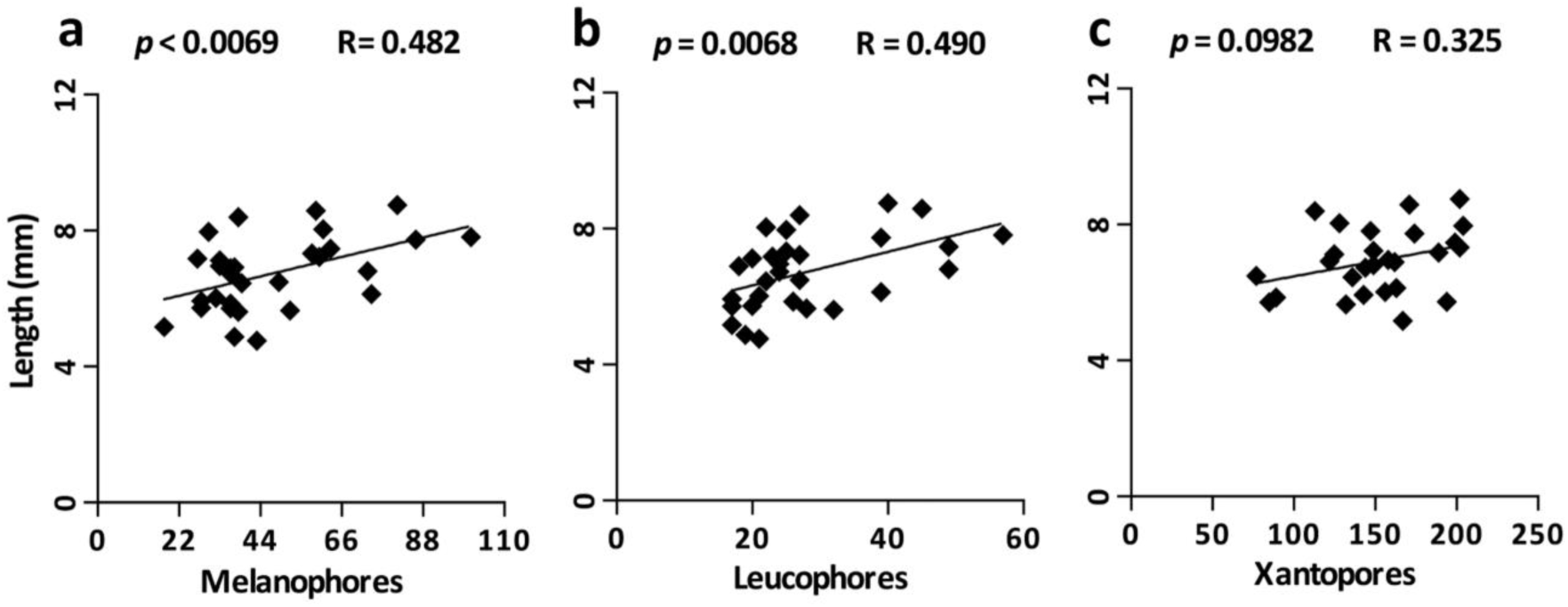
Chromatophores and body length. Pearson’s correlation coefficient between standard length and melanophores **(a)**, leucophores **(b)**, and xanthophores **(c).**

**Table S 1:**
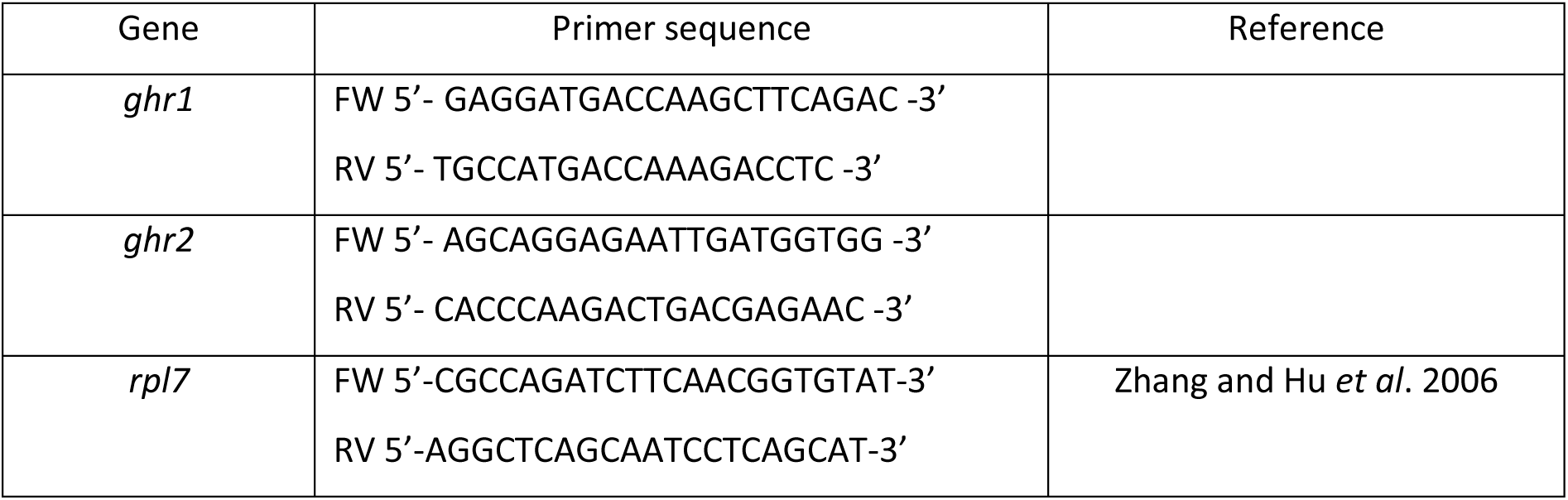
Primer sequences used in the present study for real time PCR assays.

## References

1. Fujii, R. Cytophysiology of fish chromatophores. In International Review of Cytology, 191–255 (Academic Press, 1993).

2. Sköld, H. N., Aspengren, S., Cheney, K. L., & Wallin, M.. In Fish chromatophores—from molecular motors to animal behavior. In International Review of Cell and Molecular Biology, 171–219 (Academic Press, 2016).

3. Leclercq, E., Taylor, J. F., & Migaud, H. In Morphological skin colour changes in teleosts. Fish and Fisheries. 11, 159–193 (2010).

4. Sugimoto, M. In Morphological color changes in fish: regulation of pigment cell density and morphology. Microscopy research and technique. 58, 496–503 (2002).

5. Aspengren, S., Hedberg, D., Sköld, H. N., & Wallin, M. In New insights into melanosome transport in vertebrate pigment cells. International review of cell and molecular biology. 272, 245–302 (2008).

6. Sugimoto, M., Uchida, N., & Hatayama, M. In Apoptosis in skin pigment cells of the medaka, *Oryzias latipes* (Teleostei), during long-term chromatic adaptation: the role of sympathetic innervation. Cell and tissue research. 301, 205–216 (2000).

7. Kawauchi, H., & Sower, S. A. In The dawn and evolution of hormones in the adenohypophysis. General and comparative endocrinology. 148, 3–14 (2006).

8. Benedet, S., Björnsson, B. T., Taranger, G. L., & Andersson, E. In Cloning of somatolactin alpha, beta forms and the somatolactin receptor in Atlantic salmon: seasonal expression profile in pituitary and ovary of maturing female broodstock. Reproductive biology and endocrinology. 6, 42 (2008).

9. Johnson, L. L., Norberg, B., Willis, M. L., Zebroski, H., & Swanson, P. In Isolation, characterization, and radioimmunoassay of Atlantic halibut somatolactin and plasma levels during stress and reproduction in flatfish. General and comparative endocrinology. 105, 194–209 (1997).

10. Kakisawa, S., Kaneko, T., Hasegawa, S., & Hirano, T. In Effects of feeding, fasting, background adaptation, acute stress, and exhaustive exercise on the plasma somatolactin concentrations in rainbow trout. General and comparative endocrinology. 98, 137–146 (1995).

11. Kaneko, T., & Hirano, T. In Role of prolactin and somatolactin in calcium regulation in fish. Journal of experimental biology. 184, 31–45 (1993).

12. Laiz-Carrión, R. et al. In Expression of pituitary prolactin, growth hormone and somatolactin is modified in response to different stressors (salinity, crowding and food-deprivation) in gilthead sea bream *Sparus auratus*. General and comparative endocrinology. 162, 293–300 (2009).

13. Mousa, M. A., & Mousa, S. A. In Implication of somatolactin in the regulation of sexual maturation and spawning of *Mugil cephalus*. Journal of Experimental Zoology. 287, 62–73 (2000).

14. Planas, J. V., Swanson, P., Rand-Weaver, M., & Dickhoff, W. W. In Somatolactin stimulates in vitro gonadal steroidogenesis in coho salmon, *Oncorhynchus kisutch*. General and comparative endocrinology. 87, 1–5 (1992).

15. Uchida, K. et al. cDNA cloning and isolation of somatolactin in Mozambique tilapia and effects of seawater acclimation, confinement stress, and fasting on its pituitary expression. General and comparative endocrinology. 161, 162–170 (2009).

16. Vargas-Chacoff, L. et al. Gene and protein expression for prolactin, growth hormone and somatolactin In *Sparus aurata*: seasonal variations. Comparative Biochemistry and Physiology Part B: Biochemistry and Molecular Biology. 153, 130–135 (2009)

17. Vissio, P. G. et al. Relation between the reproductive status and somatolactin cell activity in the pituitary of pejerrey, *Odontesthes bonariensis* (Atheriniformes). Journal of Experimental Zoology. 293, 492–499 (2002).

18. Cánepa, M. M., Pandolfi, M., Maggese, M. C., & Vissio, P. G. Involvement of somatolactin in background adaptation of the cichlid fish *Cichlasoma dimerus*. Journal of Experimental Zoology Part A: Comparative Experimental Biology. 305, 410–419 (2006).

19. Zhu, Y., Yoshiura, Y., Kikuchi, K., Aida, K., & Thomas, P. Cloning and phylogenetic relationship of red drum somatolactin cDNA and effects of light on pituitary somatolactin mRNA expression. General and comparative endocrinology. 113, 69–79 (1999).

20. Cánepa, M. M., Zhu, Y., Fossati, M., Stiller, J. W., & Vissio, P. G. Cloning, phylogenetic analysis and expression of somatolactin and its receptor In *Cichlasoma dimerus*: Their role in long-term background color acclimation. General and comparative endocrinology. 176, 52–61 (2012).

21. Fukamachi, S., Sugimoto, M., Mitani, H., & Shima, A. Somatolactin selectively regulates proliferation and morphogenesis of neural-crest derived pigment cells in medaka. Proceedings of the National Academy of Sciences. 101, 10661–10666 (2004).

22. Fukamachi, S., Yada, T., Meyer, A., & Kinoshita, M. Effects of constitutive expression of somatolactin alpha on skin pigmentation in medaka. Gene. 442, 81–87 (2009).

23. Fukada, H. et al. Identification of the salmon somatolactin receptor, a new member of the cytokine receptor family. Endocrinology. 146, 2354–2361 (2005).

24. Fukada, H. et al. Salmon growth hormone receptor: molecular cloning, ligand specificity, and response to fasting. General and Comparative Endocrinology. 139, 61–71 (2004).

25. Fukamachi, S., & Meyer, A. Evolution of receptors for growth hormone and somatolactin in fish and land vertebrates: lessons from the lungfish and sturgeon orthologues. Journal of molecular evolution. 65, 359–372 (2007).

26. Chen, M., Huang, X., Yuen, D. S., & Cheng, C. H. A study on the functional interaction between the GH/PRL family of polypeptides with their receptors in zebrafish: evidence against GHR1 being the receptor for somatolactin. Molecular and cellular endocrinology. 337, 114–121 (2011).

27. Pandolfi, M., Paz, D. A., Maggese, C., Ravaglia, M., & Vissio, P. Ontogeny of immunoreactive somatolactin, prolactin and growth hormone secretory cells in the developing pituitary gland of *Cichlasoma dimerus* (Teleostei, Perciformes). Anatomy and Embryology. 203, 461–468 (2001).

28. Pandolfi, M. et al. Melanin-concentrating hormone system in the brain and skin of the cichlid fish *Cichlasoma dimerus*: anatomical localization, ontogeny and distribution in comparison to α-melanocyte-stimulating hormone-expressing cells. Cell and tissue research. 311, 61–69 (2003).

29. Zhu, Y., & Thomas, P. Effects of somatolactin on melanosome aggregation in the melanophores of red drum (*Sciaenops ocellatus*) scales. General and comparative endocrinology. 105, 127–133 (1997).

30. Suzuki, M., Bennett, P., Levy, A., & Baker, B. I. Expression of MCH and POMC genes in rainbow trout (*Oncorhynchus mykiss*) during ontogeny and in response to early physiological challenges. General and comparative endocrinology. 107, 341–350 (1997).

31. Fukamachi, S., Yada, T., & Mitani, H. Medaka receptors for somatolactin and growth hormone: phylogenetic paradox among fish growth hormone receptors. Genetics. 171, 1875–1883 (2005).

32. Komine, R. et al. Transgenic medaka that overexpress growth hormone have a skin color that does not indicate the activation or inhibition of somatolactin-α signal. Gene. 584, 38–46 (2016).

33. Jeay, S., Sonenshein, G. E., Postel-Vinay, M. C., Kelly, P. A., & Baixeras, E. Growth hormone can act as a cytokine controlling survival and proliferation of immune cells: new insights into signaling pathways. Molecular and cellular endocrinology. 188, 1–7 (2002).

34. Kinoshita, M., Murata, K., Naruse K., & Tanaka, M. Medaka: Biology, Management, and Experimental Protocols. (Wiley-Blackwell, 2012).

35. Ansai, S., & Kinoshita, M. Targeted mutagenesis using CRISPR/Cas system in medaka. Biology open. 3, 362–371 (2014).

36. Castañeda Cortés, D. C., Padilla, L. F. A., Langlois, V. S., Somoza, G. M., & Fernandino, J. I. The central nervous system acts as a transducer of stress-induced masculinization through corticotropin-releasing hormone B. Development. 146, dev172866 (2019).

37. Stemmer, M., Thumberger, T., del Sol Keyer, M., Wittbrodt, J., & Mateo, J. L. CCTop: an intuitive, flexible and reliable CRISPR/Cas9 target prediction tool. PloS one. 10, e0124633 (2015).

38. Hwang, W. Y. et al. Heritable and precise zebrafish genome editing using a CRISPR-Cas system. PloS one. 8, e68708 (2013).

39. Ota, S. et al. Efficient identification of TALEN-mediated genome modifications using heteroduplex mobility assays. Genes to Cells. 18, 450–458 (2013).

40. Kinoshita, M., Kani, S., Ozato, K., & Wakamatsu, Y. Activity of the medaka translation elongation factor 1α-A promoter examined using the GFP gene as a reporter. Development, growth & differentiation. 42, 469–478 (2000).

41. Aljanabi, S. M., & Martinez, I. Universal and rapid salt-extraction of high quality genomic DNA for PCR-based techniques. Nucleic acids research. 25, 4692–4693 (1997).

42. Ramakers, C., Ruijter, J. M., Deprez, R. H. L., & Moorman, A. F. Assumption-free analysis of quantitative real-time polymerase chain reaction (PCR) data. Neuroscience letters. 339, 62–66 (2003).

43. Ruijter, J. M. et al. Amplification efficiency: linking baseline and bias in the analysis of quantitative PCR data. Nucleic acids research. 37, e45–e45 (2009).

44. Pfaffl, M. W. A new mathematical model for relative quantification in real-time RT– PCR. Nucleic acids research. 29, e45–e45 (2001).

45. Zhang, Z., & Hu, J. Development and validation of endogenous reference genes for expression profiling of medaka (*Oryzias latipes*) exposed to endocrine disrupting chemicals by quantitative real-time RT-PCR. Toxicological Sciences. 95, 356–368 (2006).

